# The evolutionarily stable distribution of fitness effects

**DOI:** 10.1101/013052

**Authors:** Daniel P. Rice, Benjamin H. Good, Michael M. Desai

## Abstract

The distribution of fitness effects of new mutations (the DFE) is a key parameter in determining the course of evolution. This fact has motivated extensive efforts to measure the DFE or to predict it from first principles. However, just as the DFE determines the course of evolution, the evolutionary process itself constrains the DFE. Here, we analyze a simple model of genome evolution in a constant environment in which natural selection drives the population toward a dynamic steady state where beneficial and deleterious substitutions balance. The distribution of fitness effects at this steady state is stable under further evolution, and provides a natural null expectation for the DFE in a population that has evolved in a constant environment for a long time. We calculate how the shape of the evolutionarily stable DFE depends on the underlying population genetic parameters. We show that, in the absence of epistasis, the ratio of beneficial to deleterious mutations of a given fitness effect obeys a simple relationship independent of population genetic details. Finally, we analyze how the stable DFE changes in the presence of a simple form of diminishing returns epistasis.

## INTRODUCTION

Mutations are the ultimate source of evolutionary change. Consequently, the distribution of their fitness effects (the DFE) is a key parameter determining the course of evolution. The DFE of new mutations controls the rate of adaptation to a new environment (Gerrish and Lenski, 1998; Good et al., 2012), the genetic architecture of complex traits (Eyre-Walker, 2010), and the expected patterns of genetic diversity and divergence (Sawyer and Hartl, 1992). To predict any of these quantities, we must first understand the shape of the DFE.

Many attempts have been made to measure the DFE or predict it from biological principles (Eyre-Walker and Keightley, 2007). Some studies have sampled directly from the DFE by measuring the fitnesses of independently evolved lines (Burch et al., 2007; Schoustra et al., 2009; Zeyl and DeVisser, 2001) or libraries of mutant genotypes (Kassen and Bataillon, 2006; McDonald et al., 2011; Sanjuán et al., 2004; Wloch et al., 2001). In other experiments, the fates of tracked lineages provide information about the scale and shape of the DFE (Frenkel et al., 2014; Imhof and Schlötterer, 2001; Perfeito et al., 2007; Rozen et al., 2002). In natural populations, the DFE leaves a signature in patterns of molecular diversity and divergence, which may be used for inference (reviewed in Keightley and Eyre–Walker 2010). A separate body of work attempts to derive the DFE from simple biophysical models of RNA (Cowperthwaite et al., 2005) or protein (Wylie and Shakhnovich, 2011).

Although these experimental and biophysical approaches can provide some insight into the shape of the DFE, they are necessarily specific to a particular organism in a particular environment. In principle, the effects of mutations depend on many biological details which vary from system to system, and it is not clear whether any general predictions are possible. However, all organisms have one thing in common: they are shaped by the process of evolution. While other phenotypes are under different selective pressures in different organisms and environments, fitness is the common currency of natural selection. It is therefore interesting to ask whether we should expect evolution to produce distributions of fitness effects with a predictable shape.

One well-known attempt to predict the shape of the DFE from evolutionary principles is the extreme value theory argument of Gillespie and Orr (Gillespie, 1983, 1984, 1991; Orr, 2003). This framework assumes that a well-adapted organism is likely to have one of the fittest available genotypes and that the fitnesses of neighboring genotypes are drawn independently from a common distribution. Gillespie and Orr argued that, under these circumstances, the fitness effects of beneficial mutations will follow an exponential distribution (provided the overall distribution of genotype fitnesses satisfies some technical conditions). This prediction has spawned a large body of theory (reviewed in Orr 2010). However, attempts to validate the theory empirically have had mixed results (e.g. Kassen and Bataillon 2006; Rokyta et al. 2008).

A limitation of extreme value theory is that it neglects the evolutionary process that produced the current genotype. Instead, it assumes that the high-fitness genotype is chosen randomly from among genotypes with similar fitness. However, different high-fitness genotypes may have very different mutational neighborhoods, and evolution does not select among these at random. Rather, it will tend to be biased toward regions of genotype space with particular properties, generating high-fitness genotypes with non-random mutational neighborhoods. This bias can lead to DFEs that are not well-characterized by extreme value theory.

Here, we use an explicit evolutionary model to study how natural selection shapes the DFE in a constant environment. When a population first encounters a given environment, it will either adapt by accumulating beneficial mutations or decline fitness in the face of an excess of deleterious mutations (Muller’s ratchet). As a population increases intness, opportunities for fitness improvement are converted to chances for deleterious back-mutation, and the fraction of mutations that are beneficial declines. Conversely, if a population declines fitness, the fraction of mutations that are beneficial increases. Eventually, the opposing forces of natural selection and Muller’s ratchet balance, and population reaches an equilibrium in which fitness neither increases nor decreases on average (Comeron and Kreitman, 2002; Goyal et al., 2012; McVean and Charlesworth, 2000; Seger et al., 2010). The approach to this equilibrium has been observed in laboratory populations (Silander et.al., 2007).)

Within this evolutionary equilibrium, a population will eventually attain a more detailed steady state. For mutations of a given absolute effect, the relative rates of beneficial and deleterious mutations will evolve until the rate of beneficial substitution exactly equals the rate of deleterious substitution (McCandlish et al., 2014; Schiffels et al., 2011). This balance holds for every effect size and therefore defines a distribution of fitness effects that is stable under the evolutionary process. This distribution serves as a natural null model for the DFE in a “well-adapted” population.

Below, we describe how the shape of the equilibrium DFE depends on the population genetic parameters and the strength of epistatic interactions across the genome. We find that, in the absence of epistasis, the equilibrium DFE has a particularly simple form and that all of the population genetic details may be summarized by a single parameter. Surprisingly, this result holds across regimes featuring very different mutational dynamics, ranging from the weak-mutation case where the equilibrium state is given by Wright’s single-locus mutation-selection-drift balance (Wright, 1931) to situations where linked selection is widespread and Wright’s results do not apply (Comeron and Kreitman, 2002; McVean and Charlesworth, 2000). We then show how epistasis changes both the shape of the equilibrium DFE and its dependence on the population genetic process.

## MODEL

We model a population of *N* haploid individuals with with an *L*-site genome, a per-genome per-generation mutation rate *U*, and a per-genome per-generation recombination rate *R*. Each site has two alleles, one conferring a fitness benefit relative to the other. The fitness difference, |*s*|, between the two alleles at each site is drawn independently from an underlying distribution *ρ*_0_ (|*s*|) with mean *s*_0_. We initially assume no epistatic interactions among sites: the fitness effect of each site is independent of the allelic state of all other sites. This simplest case functions as a null model against which deviations due to epistasis may be compared. In a later section, we expand the model to include epistasis.

**FIG. 1.**
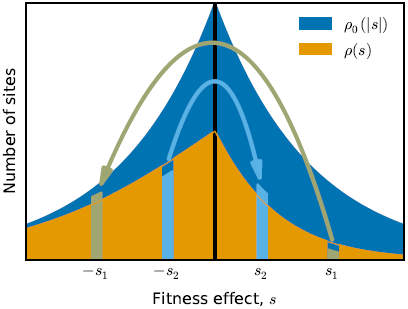
The distribution of fitness effects of mutations, *ρ*(*s*), is the product of the distribution of absolute effect sizes, *ρ*_0_(|*s*|), and the allelic state of each site. A mutation changes the DFE by changing the allelic state at that site. For example, beneficial mutation with effect +*s*_1_ creates a potential back-mutation with effect *s*_1_. Likewise, a deleterious mutation with effect *s*_2_ becomes the site of potential beneficial mutation with effect +*s*_2_.

In this model, the distribution of fitness effects, *ρ*(*s*), is determined by the distribution of absolute effects, *ρ*_0_ (|*s*|), and the genotypic state (Fig. 1). Sites carrying the deleterious allele have the potential to experience a beneficial mutation and, thus, contribute to the positive side of the DFE. Conversely, sites carrying the beneficial allele contribute to the negative side. We can therefore write the DFE as a sum of delta functions:

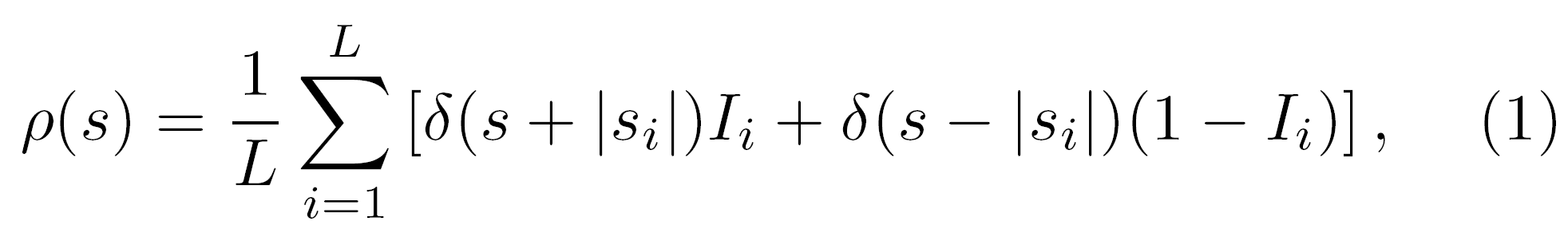

where |*s*_i_| is the absolute effect at site *i* and *I*_*i*_ = 1 if site *i* carries the beneficial allele and *I*_*i*_ = 0 otherwise. Every mutation modifies the DFE slightly by changing the allelic state at one site, removing the focal mutation from the DFE while creating the opportunity for a back-mutation with opposite effect. As the population evolves, the DFE changes until the rate of beneficial substitutions equals the rate of deleterious substitutions at every site. At this steady state, the mean change in fitness is zero and the average distribution of fitness effects is constant.

An example of this steady state, generated by a Wright-Fisher simulation of our model, is shown in Fig. 2. As observed by Seger et al. (2010), the equilibrium state is not static. Instead, the mean fitness of the population fluctuates over long timescales (Fig. 2A, orange curve) due to the cumulative fitness effect of multiple beneficial and deleterious substitutions (Fig. 2A in blue, Fig. 2B). Consistent with the steady state assumption, the fitness effects of fixed mutations are roughly symmetric about zero (Fig. 2C), with deviations due to the relatively small size of the sample shown here. The magnitudes of these fixed fitness effects reflect the population dynamics as well as the shape of the equilibrium DFE; in this example, they are at most of order 10/*N*. We will discuss the relationship between the fitness effects of fixed mutations and the population parameters below.

Our example simulation also illustrates complications that arise due to linked selection. For example, in Fig. 2B, we see that mutations mostly fix in clusters. The phenomenon of clustered fixations is a signature of linked selection that has been predicted in theory (Park and Krug, 2007) and observed in experimental and natural populations (Lang et al., 2013; Nik-Zinal et al., 2012; Strelkowa and Lässig, 2012). In Fig. 2D we also see that linked selection reduces the fixation probability of beneficial mutations relative to the standard single-locus prediction (Wright, 1931) in a way that cannot be summarized by a simple reduction in the effective population size.

Finally, our simulations reveal the equilibrium shape of the DFE (Fig. 2E). In this example, the underlying distribution of absolute effects, *ρ*_0_ |*s*|, is exponential with mean *Ns*_0_ = 10. However, the equilibrium distribution of beneficial effects, ρ(s), falls off much faster. In fact, almost no beneficial mutations are available with effects greater than *Ns* = 10.

## ANALYSIS

### The stable distribution of fitness effects

To obtain analytical expressions for the steady state DFE in our model, we focus on the large *L* limit, where we can neglect differences in the DFE between genotypes that segregate simultaneously in the population. With this assumption, the average DFE evolves according to the differential equation

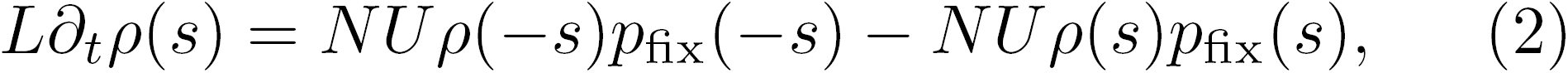

where *p*_fix_(*s*) is the fixation probability of a mutation with effect *s*. The first term on the right-hand-side of Eq. (2) represents the substitution rate of deleterious mutations with absolute effect |*s*|, which is the product of the mutation supply rate and the fixation probability. Likewise, the second term gives the substitution rate of beneficial mutations with effects. Eq. (2) captures the fact that each mutation changes the DFE slightly by converting a beneficial mutation to a potential deleterious mutation, or vice versa (Fig. 1).

At long times, the DFE evolves toward the equilibrium state *ρ*_eq_(*s*), in which the substitution rates of beneficial and deleterious mutations exactly balance for every value of *s*. Setting the time derivative in Eq. (2) to zero yields the equilibrium DFE ratio:

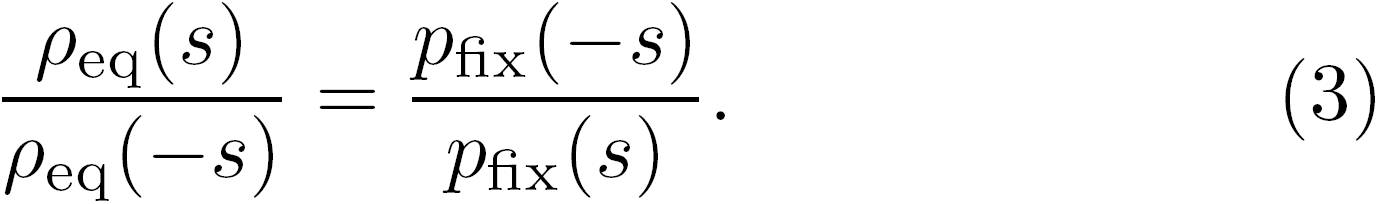

We can rewrite Equation (3) in terms of the underlying distribution of absolute effects and the equilibrium state of the genome:

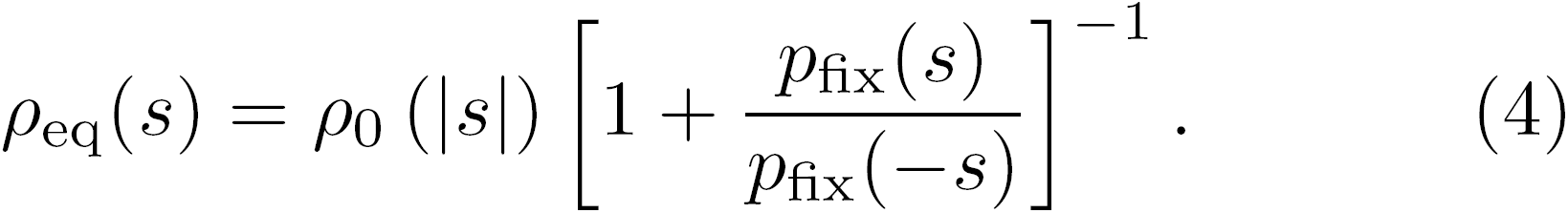

Eq. (4) shows that the equilibrium DFE is determined by the relative probabilities of fixation of beneficial and deleterious mutations. Unfortunately, there is no general expression for these fixation probabilities because they depend on the effects of linked selection (Hill and Robertson, 1966). Moreover, these dynamics of linked selection depend on the shape of the DFE, so the right hand side of Eq. (4) implicitly depends on *ρ*_eq_(*s*) in.

Fortunately, there are two limits of our model where simple expressions for *p*_fix_(*s*) are available. In the limit that mutations are rare (*NU*→ 0), each mutation fixes or goes extinct independently. Thus, we can use the single-locus probability of fixation (Fisher, 1930; Wright, 1931),

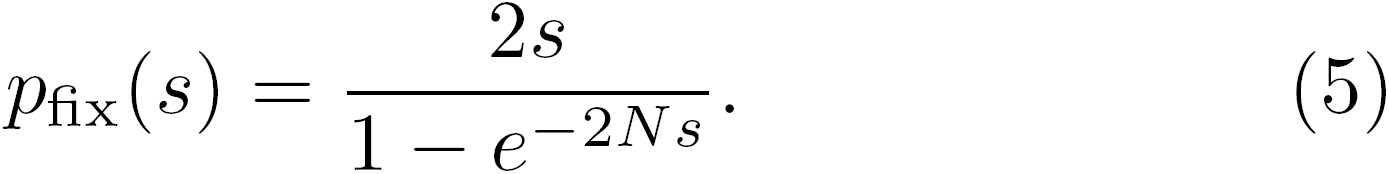

Substituting Eq. (5) into Eq. (4) yields

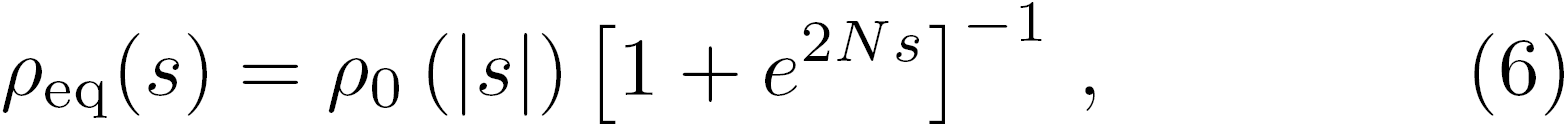

which is the familiar single-locus mutation-selection-drift equilibrium (Wright, 1931).

In the opposite extreme, where the mutation rate is very high, previous work has shown that the probability of fixation depends exponentially on *s*:

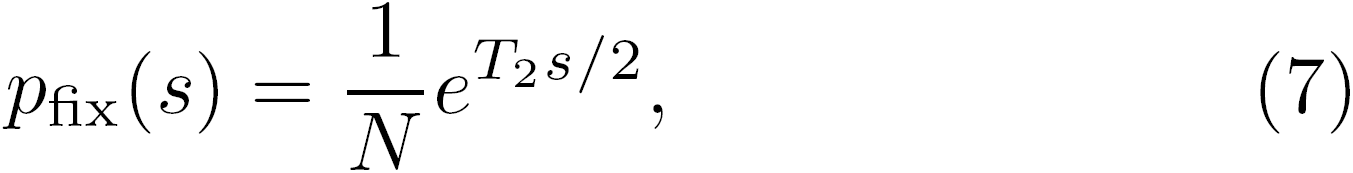

where *T*_2_ is the average pairwise coalescence time (Good et al., 2014; Neher et al., 2013). Note that linked selection alters the functional form of *p*_fix_(*s*), and hence cannot be captured by a simple reduction in effective population size. In this strong mutation limit, substituting Eq. (7) into Eq. (4) shows that the equilibrium DFE has the form:

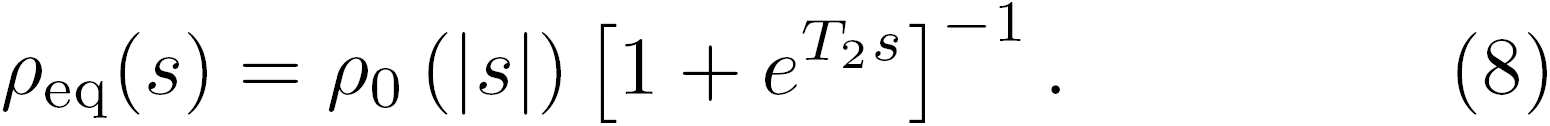

Surprisingly, the shape of the equilibrium DFE has the same dependence on s in both limiting regimes. This is because the ratio *p*_fix_(−*s*) / *p*_fix_(*s*) falls off exponentially with *s* when mutation is weak as well as when it is very strong, even though the fixation probabilities have different forms. The fact that the DFE ratio has the same form in two very different limiting regimes suggests that the result may be general. We therefore propose that 

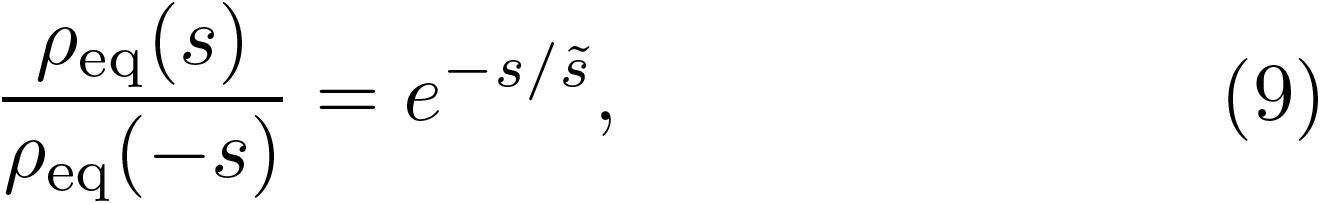

where 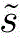
is the scale at which the DFE ratio falls off with *s*. This single scale encapsulates all of the effects of linked selection and their dependence on the underlying parameters. In the weak mutation limit, 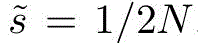 while in the strong mutation limit, 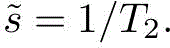

**FIG. 2.**
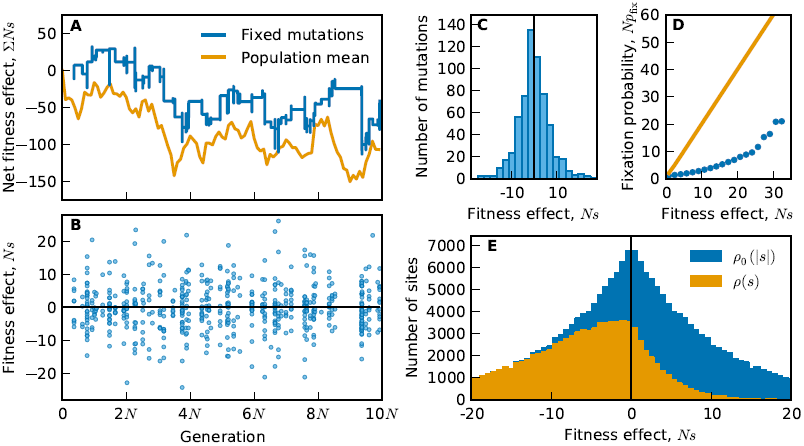
An example of the steady state dynamics, generated by a Wright-Fisher simulation of our model (*N* = 10^4^, *NU* = 10^2^, *NR* = 0, *ρ*_0_ (|*s*|) ~ Exp[10/*N*], *L* = 10^5^). A) Time course of the mean fitness of the population and the cumulative effect of fixed mutations. B) The fitness effect of each fixed mutation versus its fixation time. C) Histogram of the fitness effects of all fixed mutations. D) The fixation probability of a beneficial mutation as a function of its fitness effect. Simulation results are shown in circles; the single-locus theory prediction in the absence of linked selection is shown as the solid line. E) The distribution of absolute effects *ρ*_0_(|*s*|) (blue) and distribution of fitness effects *ρ*(*s*) (orange) measured at the end of the simulation.

To test this conjecture, we calculated the DFE ratio, 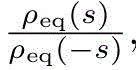 across a broad range of parameters for an asexual population with exponential *ρ*_0_(|*s*|). For each set of parameters, we found the evolutionary equilibrium by varying the initial fraction of sites fixed for the deleterious allele and recording the fitness change in the simulation. We varied the length of the simulations in order to verify convergence to the steady state (9704File S1, Fig. S1). As predicted, we found that the DFE ratio declines exponentially with *s* for all population parameters (Fig. 3A). Similar results are obtained for other choices of *ρ*_0_(|*s*|) and in the presence of recombination (Fig. S2). The observed values of 2*N*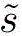varied over three orders of magnitude for the parameters tested (Fig. 3A inset). When mutation is weak, 2*N* 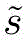 ≈ 1, in accordance with the single-locus intuition. Figure 3A confirms that the limiting analysis above is general: for the purpose of determining the equilibrium DFE, the net result of the complicated mutational dynamics can be summarized by the single parameter 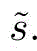

### The steady state substitution rate

While the form of the equilibrium DFE is independent of the mutational dynamics, other features of the steady state depend in detail on the extent of linked selection. For example, as shown in Fig. 2A-C, the steady state is characterized by a constant “churn” of fixations. The distribution of fitness effects of the mutations that fix is symmetrical and its shape is determined by the substitution rate *K* as a function of |*s*|. In order to compare across simulations with different overall mutation rates and underlying DFEs, we define a normalized substitution rate by dividing *K*(|*s*|) by the rate at which mutations of effect |*s*| arise. Using Eq. (5) and Eq. (7), we can make analytical predictions about the substitution rate in both the weak and strong mutation limits. We find that

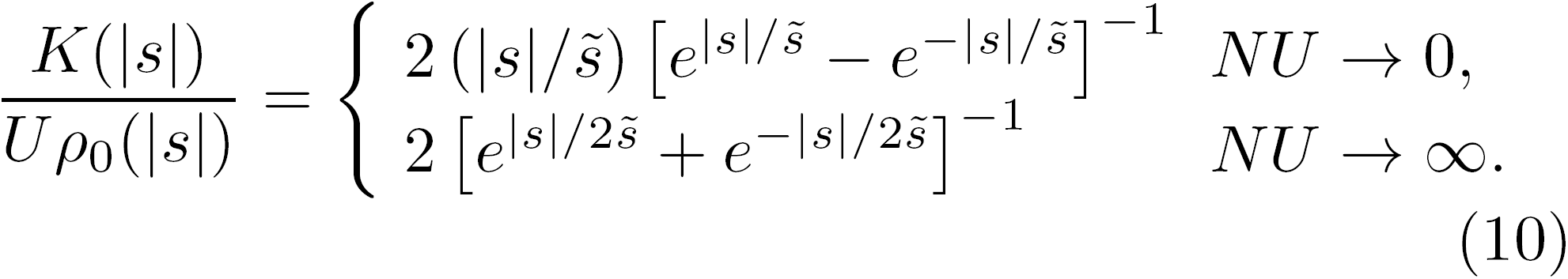

Note that *K*(|*s*|) has a different functional form in the two limits.

**FIG. 3.**
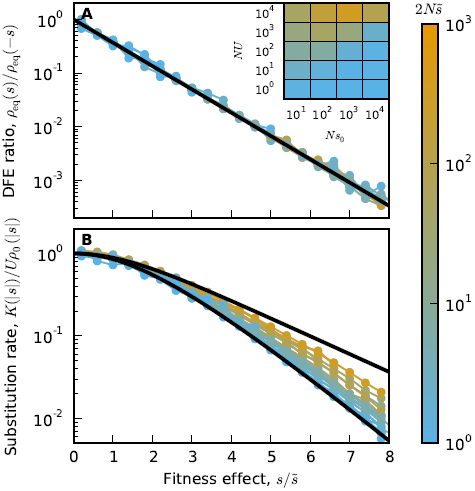
The equilibrium DFE and steady-state substitution rate. A) The equilibrium ratio of the beneficial mutation rate to the deleterious mutation rate for mutations with absolute effect |*s*|, averaged over 100 replicate simulations. Fitness effects are scaled by a parameter 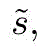 fit to the average final DFE for each parameter set. The color of each line indicates the value of 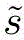 for each parameter set (inset). The solid black line shows exp(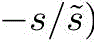. B) The substitution rate of mutations with absolute effect |*s*|, *K*(|*s*|), declines with the scaled fitness effect 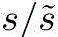 Upper and lower solid curves give the analytical results for the strong and weak-mutation limits respectively. All results here are for an asexual population with exponential *ρ*_0_ (|*s*|). The results for other choices of *ρ*_0_ (|*s*|) and for *R* > 0 are equivalent (Fig. S2).

Fig. 3B shows the observed substitution rates in our simulations as a function of the scaled fitness effect 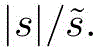 Here, the values of the scaling parameter 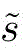 are the values t to the equilibrium DFE for each parameter set. The two limiting predictions from Eq. (10) are shown as solid curves. These predictions bracket the observed substitution rates. As expected, when 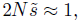, the substitution rates approach the weak mutation limit. On the other hand, when 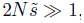, there is a higher rate of substitution for each value 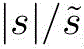 approaching but not achieving the strong mutation limit.

The relationship between the substitution rate and the effect of the mutation has two notable features. First, the substitution rate declines with the fitness effect, because at equilibrium large-effect sites are almost always fixed for the beneficial allele. Second, unlike the DFE ratio, the substitution rate is not a function of the scaled parameter 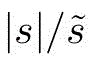 alone. Instead, populations with large 2*N* 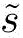 tend to have higher substitution rates of mutations with any given effect than populations with small 2*N* 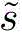 due to the effects of linked selection.

### The coalescent timescale determines the equilibrium DFE

So far we have treated 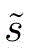 as a fitting parameter, but we will now argue that it can be interpreted in terms of a fundamental timescale of the evolutionary process. Eq. (9) shows that 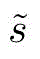 is the scale at which a mutation transitions from being effectively neutral to experiencing the effects of selection. This scale is set by the coalescent timescale on which the future common ancestor of the population is determined (Good and Desai, 2014). For example, a deleterious mutation with cost *s* is typically purged from the population in *s* ^−1^ generations. If *s* ^−1^ is much shorter than the time it takes to choose a future common ancestor, the mutant lineage will be eliminated before it has an opportunity to fix. On the other hand, if *s* ^−1^ is much larger than the coalescent timescale, selection will not have enough time to influence the fate of the mutant. We therefore expect 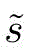 to be of order the inverse of the coalescent timescale.

The coalescent timescale depends on the complicated interplay between drift, selection, and interference. Thus, it is di cult to predict from the underlying parameters. Furthermore, the coalescent timescale and the DFE depend on one another and change together as the population evolves. Fortunately, this timescale also determines an independent quantity: the level of neutral diversity within the population. Therefore, we should be able to predict 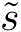 from measurements of diversity in our simulated populations.

To test this expectation, we introduced neutral mutations into our equilibrium simulations and measured the average number of pairwise differences, *π*, normalized by the expected diversity in a neutrally evolving population of the same size (*π*_0_ = 2*NU*). As expected, Fig. 4 shows that 2*N* 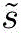 is inversely proportional to π/π_0_. Furthermore, the observed relationship interpolates between the strong-mutation prediction 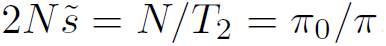 and the weak-mutation prediction 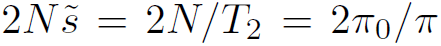. Thus, we can predict the fitted DFE ratio parameter from the neutral pairwise diversity up to an 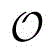 (1) constant.

**FIG. 4.**
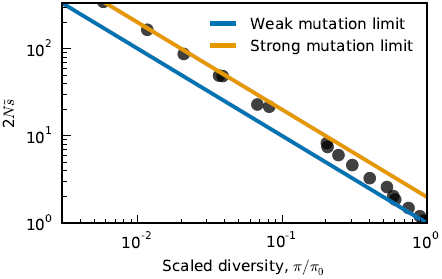
Pairwise diversity predicts the equilibrium DFE ratio. For each parameter set, we t the scaling parameter 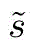 to the equilibrium DFE and measured the average number of pairwise differences, *π*, normalized by the expected diversity in a neutrally evolving population of the same size, *π*_0_ = 2*NU*.

### Diminishing returns epistasis

In the previous sections, we have considered a model without epistasis, where the fitness effect of each site is independent of the state of all other sites. While this provides a useful null model, it is interesting to consider the effect of epistatic interactions on the equilibrium distribution of fitness effects. There are many possible models of epistasis that we could consider. Here, we focus on a simple example suggested by recent empirical work: a general pattern of diminishing returns epistasis (Chou et al., 2011; Khan et al., 2011; Kryazhimskiy et al., 2014; Wiser et al., 2013).

In the simplest case, this type of epistasis arises when fitness is a nonlinear function of a phenotypic trait. Here, the fitness effect of a mutation is not fixed, but depends on the state of the genome through the current phenotypic value. Specifically, we consider a single fitness-determining phenotypic trait, *ξ* controlled by *L* additive sites:

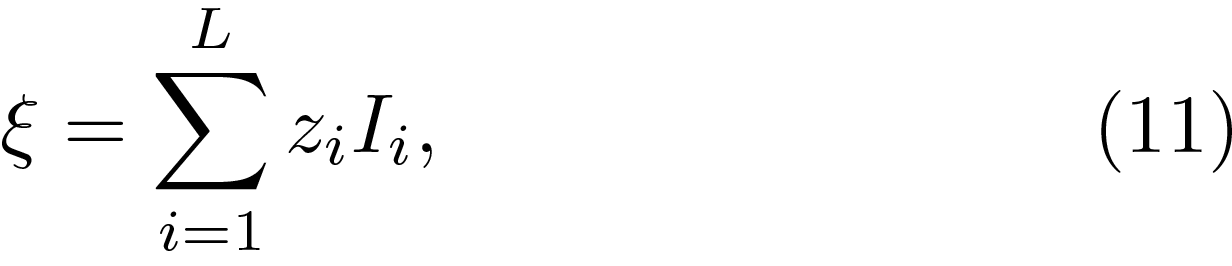

where *z*_*i*_ is the phenotypic effect of site *i* and *I*_*i*_ ∈ {0, 1} is an indicator variable denoting the allelic state at that site. The fitness of an individual with phenotype *ξ* is then given by some function *F*(*ξ*).

In this epistatic case, Eq. (2) no longer applies because the fitness effect of a mutation depends on the current value of the phenotype. However, because we assume that mutations interact additively at the level of phenotype, we can write an analogous equation for the distribution of phenotypic effects, *φ*(*z*):

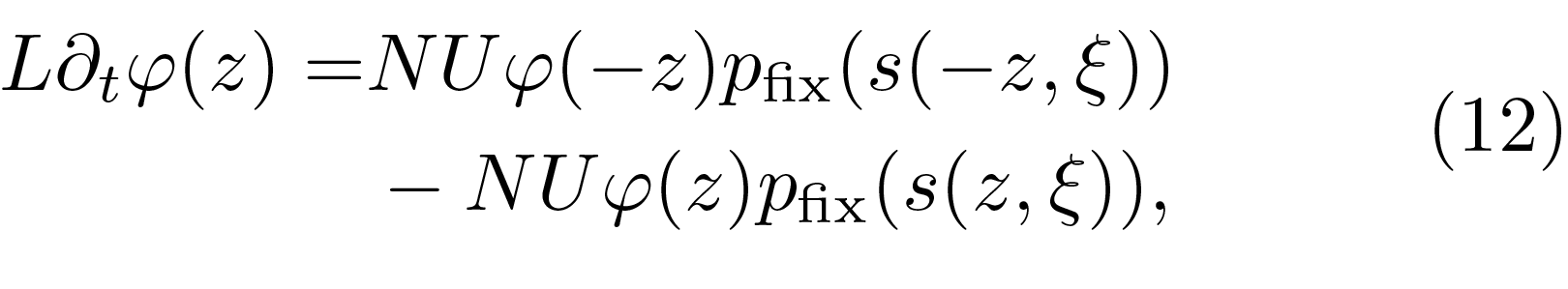

where *s*(*z*, *ξ*) = *F*(ξ + *z*) − *F*(*ξ*) is the fitness effect of a mutation of phenotypic effect *z* that occurs in an individual with phenotype *ξ*. By analogy to Eq. (4), the equilibrium distribution of phenotypic effects, *φ*_*eq*_(*z*), is then given by

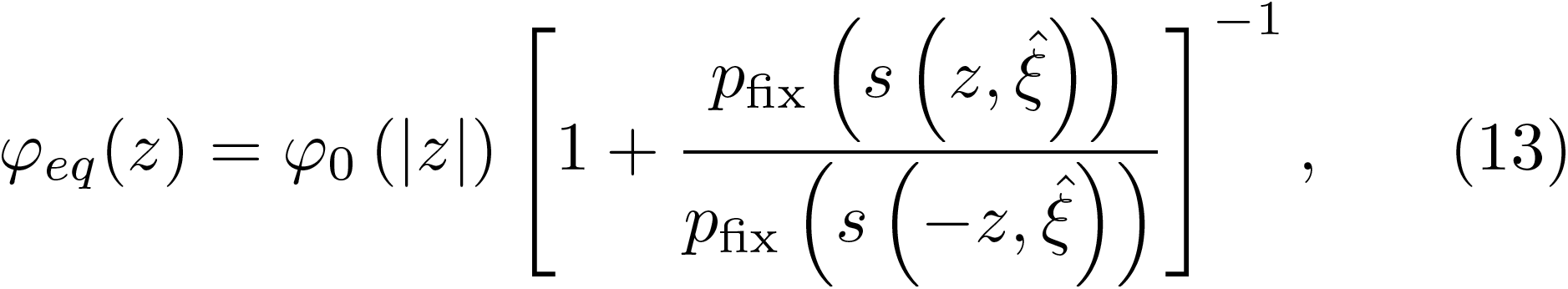

where *φ*_0_(|*z*|) is the distribution of absolute phenotypic effects and 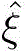 is the equilibrium phenotypic value, which depends on the strength of epistasis. In order to find the equilibrium distribution of fitness effects, we must change variables from phenotypic effect to fitness effect:

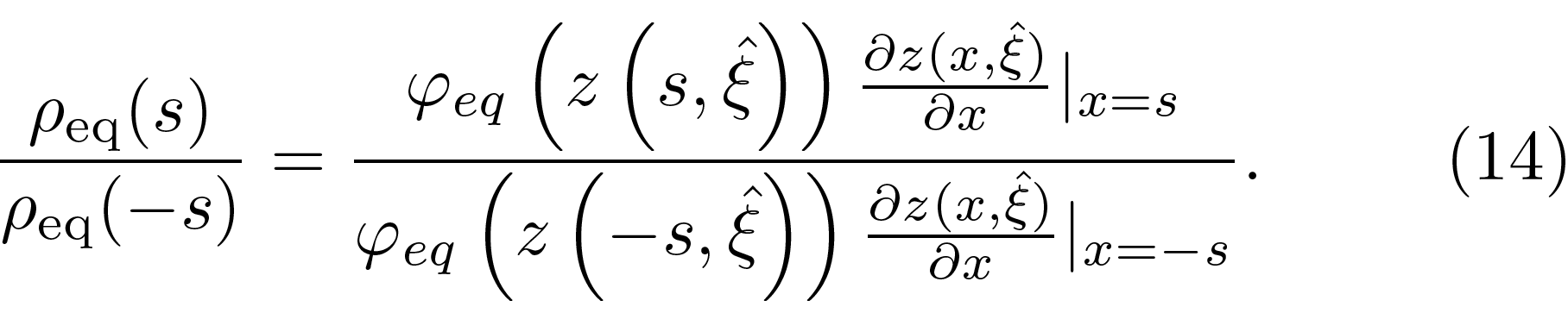

Although Eq. (14) is di cult to interpret in general, we can gain qualitative insight by considering the limit of weak diminishing returns epistasis, where we can expand *F*(*ξ*) in the form

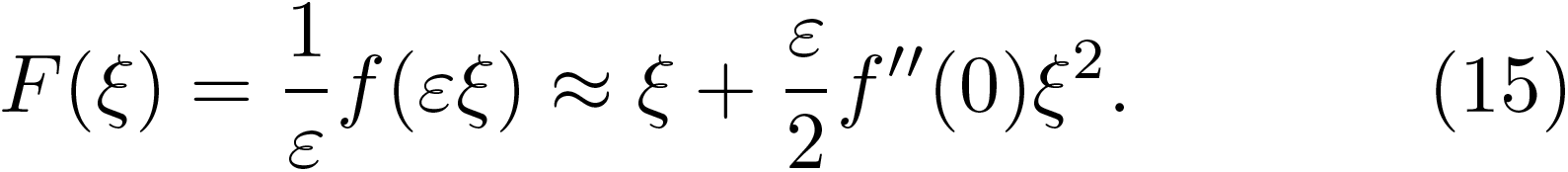

Here *ε* ≪ 1 is a constant that determines the scale at which epistatic effects become important, and *f"*(0) < 0 since we assume diminishing returns. In this limit, we have

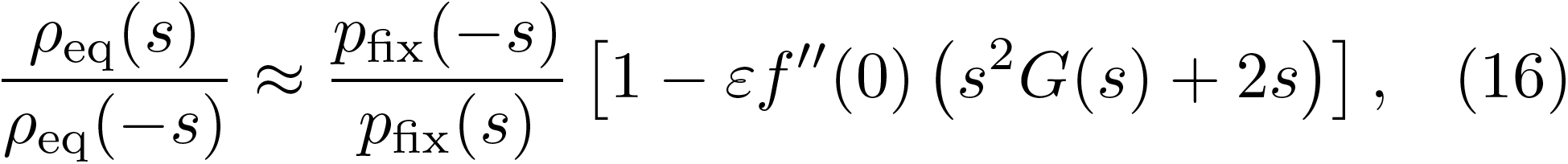

where we have defined

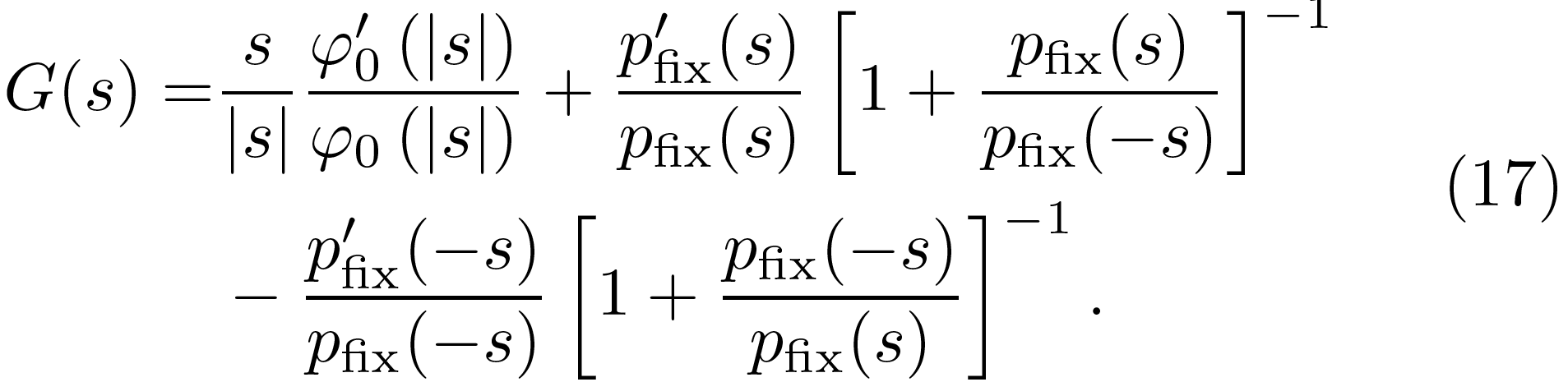

Note that epistasis introduces dependence on the log derivative of the fixation probability: *p'*_fix_(*s*) / *p*_fix_(*s*). Unlike the ratio *p*_fix_(−*s*) / *p*_fix_(*s*), this quantity does depend on the details of the evolutionary dynamics. For example, in the strong-mutation limit the log derivative is a positive constant, while in the weak-mutation limit it is a positive and increasing function of *s*. As a result, epistasis has an influence on the equilibrium distribution of fitness effects that cannot be captured by the parameter 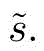

To see the effect of epistasis on the shape of the DFE more concretely, consider the case where the strong-mutation limit applies and 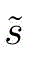 ≪ *s*_0_. Under these conditions, *G*(*s*) ~ 1/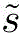 and the first order correction in Eq. (16) is positive for *s* ≪ 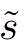 and negative for *s* ≫ 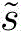. As a result, there are more weakly beneficial mutations and fewer strongly beneficial mutations available in the presence of diminishing returns epistasis than the non-epistatic analysis would predict.

## DISCUSSION

The evolutionary process shapes the distribution of available mutations. Here, we have calculated the equilibrium DFE that evolution produces in a simple null model of a finite genome with no epistasis. Across a wide range of parameters, this equilibrium DFE has the property that 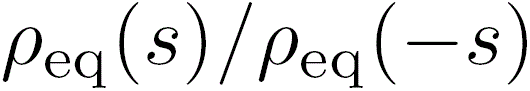 falls off exponentially with *s*. This property holds despite very different population dynamics for different parameters. It is also independent of the shape of the underlying DFE and rate of recombination. The rate of exponential decline depends on the coalescent timescale, which can be predicted from the neutral diversity in the population.

Our results for the equilibrium DFE are strikingly different from earlier attempts to deduce features of the DFE from extreme value theory arguments (Gillespie, 1983, 1984, 1991; Orr, 2003). According to extreme value theory, the DFE of a well-adapted population only depends on the distribution of genotype fitnesses, and not on the particular evolutionary history that brought the population to its well-adapted state. In contrast, we have shown here that the population genetic process (e.g., the historical population size and the mutation rate) can strongly influence the both the shape and scale of the equilibrium DFE, even when the distribution of genotype fitnesses is held constant. As an example, consider the case where *ρ*_0_ (|*s*|) is a half-normal distribution, so that the equilibrium DFE is given by

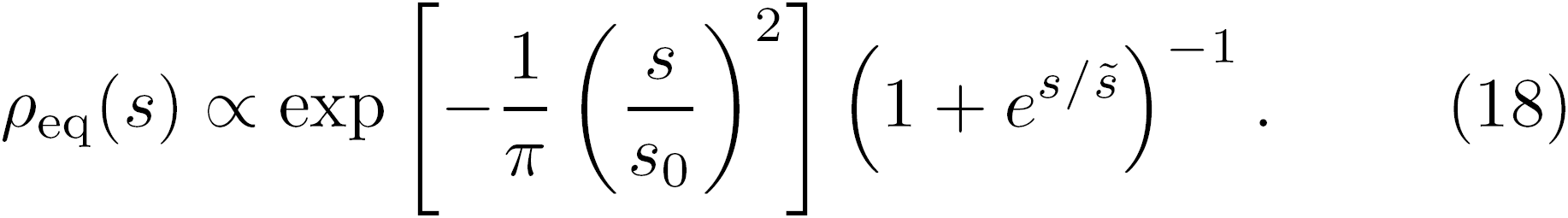

The equilibrium DFE is thus determined by two scales: s_0_, the scale of the underlying DFE, and 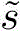 the fitness scale at which sites feel the effects of selection strongly. When *s*_0_ ≪ 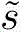 selection barely biases the allelic states and 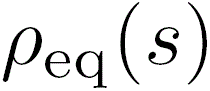 is Gaussian. Conversely, when *s*_0_ ≫ 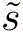 the equilibrium DFE falls o exponentially for large s. This simple example shows that the shape of the DFE can strongly depend on both the population genetic parameters and the shape of the underlying genotype distribution, and there is no reason to expect it to be exponential in general. In contrast, our analysis predicts that in the absence of epistasis the equilibrium DFE ratio 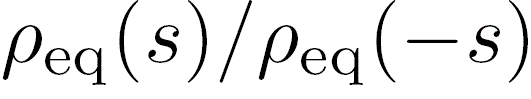 should have a simple exponential form; this can in principle be directly tested experimentally.

Our prediction for the DFE ratio has the same form as standard mutation-selection-drift balance at a single locus, where *N* is replaced by an "effective population size"

*N*_e_ = 1 / 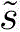 which can be estimated from neutral diversity. This drift-barrier intuition forms the basis for many previous empirical studies (Loewe and Charlesworth, 2006; Lohmueller et al., 2008; Sung et al., 2012) and theoretical work on the evolution of the mutation rate (Lynch, 2011). To some extent, the robustness of this single-locus prediction is surprising, given that it appears to hold even when sites do not evolve independently. Our analysis shows how this simple result emerges more generally and illustrates how it breaks down in the presence of epistasis. In addition, we have shown that the single-locus analysis fails to predict the substitution rate. Thus, while drift barrier arguments can correctly predict the probability a given locus is fixed for the beneficial allele, they will often substantially underestimate the flux of fixations of both beneficial and deleterious alleles, even after accounting for the reduction in e effective population size.

This constant flux of fixations emphasizes the dynamic nature of the equilibrium that we study here. Rather than approaching a static fitness peak, a population adapting to a constant environment will eventually approach this dynamic steady state. As this happens, the rate of fitness evolution slows down over time, eventually reaching a fitness plateau where mean fitness does not change on average, while rates of molecular evolution remain high. Depending on the underlying parameters, this population genetic limit to optimization can occur long before any absolute physiological limits become relevant.

In the present work, we have studied only the simplest model of the evolution of the DFE. This null model has several key limitations, which present interesting avenues for future work. Most importantly, we have focused only on evolution in a constant environment. We expect a similar dynamic equilibrium to arise in a fluctuating environment, provided that the statistics of these fluctuations remain constant through time (Gillespie, 1991; Mustonen and Lässig, 2009). To analyze this more complex situation, we need to understand the distribution of pleiotropic effects of mutations across environmental conditions, and how this pleiotropy affects fixation probabilities.

Another important limitation of our model is that we have only considered one specific form of epistasis: a general diminishing returns model suggested by recent microbial evolution experiments. This type of epistasis leads to an excess of weakly beneficial mutations relative to the non-epistatic case, in a way that crucially depends on the population genetic parameters. However, many other types of epistasis may also be common in natural populations. For example, idiosyncratic interactions between specific mutations, including sign epistasis, have been observed in several systems (de Vos et al., 2013; Weinreich et al., 2006). We also often expect to observe modular interactions, in which only the first mutation in each module can confer a fitness effect (Tenaillon et al.,2012). As more detailed empirical measurements of epistasis across the genome become available, it will be interesting to analyze how these effects change the expected steady state DFE.

Despite these limitations, our analysis provides a useful null model for how the process of evolution shapes the distribution of fitness effects. Our results suggest that experiments should seek to measure the DFE ratio, Eq. (3), which in the absence of epistasis is independent of the mutational dynamics or the underlying distribution of effects. Deviations from the null prediction may be informative about the global structure of epistasis or the evolutionary history of the population.

---

**FIG. S1.**
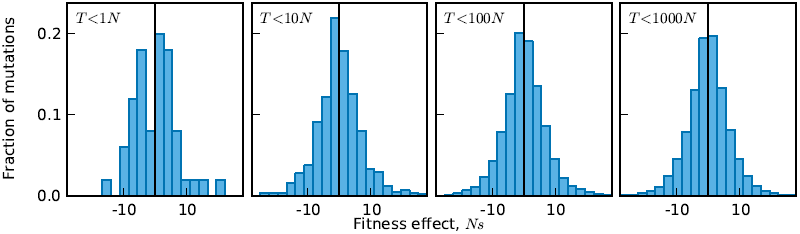
The distribution of fixed effects over time in the example simulation shown in Fig. 2. Each panel shows the effects of all mutations fixed by time *T*. The *T* < 10*N* panel corresponds to Fig. 2B. To check for convergence to the steady state, we ran each set of simulations until the distribution of fixed effects was symmetric about zero.

**FIG. S2.**
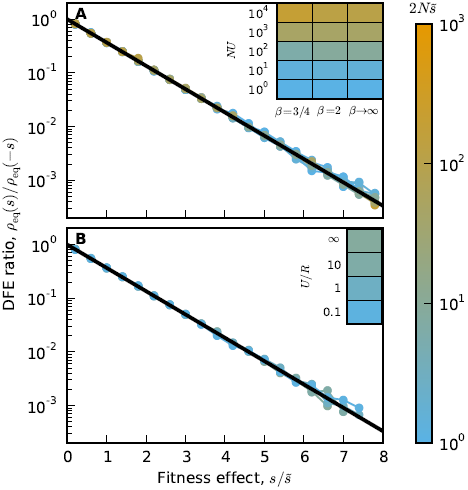
The equilibrium DFE for different underlying DFEs and recombination rates. A) The equilibrium ratio of beneficial mutations to deleterious mutations for mutations with absolute effect |*s*|, averaged over 100 replicate simulations. We examined the effect of underlying DFEs in the stretched exponential family, 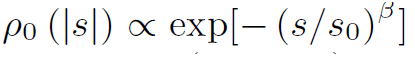. Specifically, we simulated heavy-tailed stretched exponential (β = 3/4), half-Gaussian (β = 2), and uniform (β → ∞) underlying DFEs. *Ns*_0_ = 100 for all simulations. B) The equilibrium DFE ratio in populations with recombination. As the recombination rate increases, 2*N*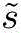 → 1 as mutations begin to fix independently. *NU* = 100, *Ns*_0_ = 10 for all simulations. We used FFPopsim (Zanini and Neher, 2012) to simulate recombining populations.

## REFERENCES

Burch, C. L., Guyader, S., Samarov, D., and Shen, H. 2007. Experimental estimate of the abundance and effects of nearly neutral mutations in the rna virus *φ*6 . Genetics 176:467–476.

Chou, H.-H., Chiu, H.-C., Delaney, N. F., Segré, D., and Marx, C. J. 2011. Diminishing returns epistasis among beneficial mutations decelerates adaptation. Science 332:1190–1192.

Comeron, J. M. and Kreitman, M. 2002. Population, evolutionary, and genomic consequences of interference selection. Genetics 161:389–410.

Cowperthwaite, M. C., Bull, J., and Meyers, L. A. 2005. Distributions of beneficial fitness effects in rna. Genetics 170:1149–1457.

de Vos, M. G. J., Poelwijk, F. J., Battich, N., Ndika, J. D. T., and Tans, S. J. 2013. Environmental dependence of genetic constraint. PLoS Genet 9:e1003580.

Eyre-Walker, A. 2010. Genetic architecture of a complex trait and its implications for fitness and genome-wide association studies. Proc Natl Acad Sci 107:1752–1756.

Eyre-Walker, A. and Keightley, P. D. 2007. The distribution of fitness effects of new mutations. Nat Rev Gen 8:610–618.

Fisher, R. A. 1930. The distribution of gene ratios for rare mutations. Proc Roy Soc Edinburgh 50:204–219.

Frenkel, E. M., Good, B. H., and Desai, M. M. 2014. The fates of mutant lineages and the distribution of fitness effects of beneficial mutations in laboratory budding yeast populations. Genetics 196:1217–1226.

Gerrish, P. and Lenski, R. 1998. The fate of competing beneficial mutations in an asexual population. Genetica 127:127–144.

Gillespie, J. 1983. A simple stochastic gene substitution model. Theoretical Population Biology 23:202–215.

Gillespie, J. 1984. Molecular evolution over the mutational landscape. Evolution 38:1116–1129.

Gillespie, J. 1991. The causes of molecular evolution. Oxford University Press, New York.

Good, B. H. and Desai, M. M. 2014. Deleterious passengers in adapting populations. Genetics 198:1183–1208.

Good, B. H., Rouzine, I. M., Balick, D. J., Hal-latschek, O., and Desai, M. M. 2012. Distribution of fixed beneficial mutations and the rate of adaptation in asexual populations. Proc. Natl. Acad. Sci. 109:4950–4955.

Good, B. H., Walczak, A. M., Neher, R. A., and Desai, M. M. 2014. Genetic diversity in the interference selection limit. PLoS Genetics 10:e1004222.

Goyal, S., Balick, D. J., Jerison, E. R., Neher, R. A., Shraiman, B. I., and Desai, M. M. 2012. Dynamic mutation-selection balance as an evolutionary attractor. Genetics 191:1309–1319.

Hill, W. G. and Robertson, A. 1966. The effect of linkage on limits to artificial selection. Genet. Res. 8:269–294.

Imhof, M. and Schlötterer, C. 2001. Fitness effects of advantageous mutations in evolving Escherichia coli populations. Proc. Natl. Acad. Sci. USA 98:1113–1117.

Kassen, R. and Bataillon, T. 2006. Distribution of t-ness effects among beneficial mutations before selection in experimental populations of bacteria. Nature Genetics 38:484–488.

Keightley, P. D. and Eyre-Walker, A. 2010. What can we learn about the distribution of fitness effects of new mutations from dna sequence data? Philosophical Transactions of the Royal Society B: Biological Sciences 365:1187–1193.

Khan, A. I., Din, D. M., Schneider, D., Lenski, R. E., and Cooper, T. F. 2011. Negative epistasis between beneficial mutations in an evolving bacterial population. Science 332:1193–1196.

Kryazhimskiy, S. K., Rice, D. P., Jerison, E. R., and Desai, M. M. 2014. Global epistasis makes adaptation predictable despite sequence-level stochasticity. Science 344:1519–1522.

Lang, G. I., Rice, D. P., Hickman, M. J., Sodergren, E., Weinstock, G. M., Botstein, D., and Desai, M. M. 2013. Pervasive genetic hitchhiking and clonal interference in forty evolving yeast populations. Nature 500:571–574.

Loewe, L. and Charlesworth, B. 2006. Inferring the distribution of mutational effects on fitness in drosophila. Biol Lett 2:426–430.

Lohmueller, K. E., Indap, A. R., Schmidt, S., Boyko, A. R., Hernandez, R. D., Hubisz, M. J., Sninsky, J. J., White, T. J., Sunyaev, S. R., Nielsen, R., et al. 2008. Proportionally more deleterious genetic variation in european than in african populations. Nature 451:994–997.

Lynch, M. 2011. The lower bound to the evolution of mutation rates. Genome Biol. Evol. 3:1107–1118.

McCandlish, D. M., Otwinowski, J., and Plotkin, J. B. 2014. On the role of epistasis in adaptation. arXiv

McDonald, M. J., Cooper, T. F., Beaumont, H. J., and Rainey, P. B. 2011. The distribution of fitness effects of new beneficial mutations in pseudomonas fluorescens. Biology letters 7:98–100.

McVean, G. A. T. and Charlesworth, B. 2000. The effects of hill-robertson interference between weakly selected mutations on patterns of molecular evolution and variation. Genetics 155:929–944.

Mustonen, V. and Lässig, M. 2009. From fitness landscapes to seascapes: non-equilibrium dynamics of selection and adaptation. Trends Genet 25:111–119.

Neher, R. A., Kessinger, T. A., and Shraiman, B. I. 2013. Coalescence and genetic diversity in sexual populations under selection. Proc Nat Acad Sci 110:15836–15841.

Nik-Zinal, S., Loo, P. V., Wedge, D. C., alexandrov, L. B., and et al 2012. The life history of 21 breat cancers. Cell 149:994–1007.

Orr, H. 2003. The distribution of fitness effects among beneficial mutations. Genetics 163:1519–1526.

Orr, H. A. 2010. The population genetics of beneficial mutations. Philosophical Transactions of the Royal Society B: Biological Sciences 365:1195–1201.

Park, S. and Krug, J. 2007. Clonal interference in large populations. Proc. Natl. Acad. Sci. USA 104:18135–18140.

Perfeito, L., Fernandes, L., Mota, C., and Gordo, I. 2007. Adaptive mutations in bacteria: High rate and small effects. Science 317:813.

Rokyta, D. R., Beisel, C. J., Joyce, P., Ferris, M. T., Burch, C. L., and Wichman, H. A. 2008. Beneficial fitness effects are not exponential for two viruses. J. Mol. Evol. 67:368–376.

Rozen, D. E., de Visser, J. A., and Gerrish, P. J. 2002. Fitness effects of beneficial mutations in microbial populations. Curr Biol 12:1040–1045.

Sanjuän, R., Moya, A., and Elena, S. F. 2004. The distribution of fitness effects caused by single-nucleotide substitutions in an rna virus. Proc. Natl. Acad. Sci. USA 101:8396–8401.

Sawyer, S. A. and Hartl, D. L. 1992. Population genetics of polymorphism and divergence. Genetics 132:1161–1176.

Schiffels, S., Szöllösi, G., Mustonen, V., and Lässig, M. 2011. Emergent neutrality in adaptive asexual evolution. Genetics 189:1361–1375.

Schoustra, S. E., Bataillon, T., Gifford, D. R., and Kassen, R. 2009. The properties of adaptive walks in evolving populations of fungus. PLoS biology 7:e1000250.

Seger, J., Smith, W. A., Perry, J. J., Hunn, J., Kaliszewska, Z. A., et al. 2010. Gene geneologies strongly distorted by weakly interfering mutations in constant environments. Genetics 184:529–545.

Silander, O. K., Tenaillon, O., and Chao, L. 2007. Understanding the evolutionary fate of finite populations: The dynamics of mutational effects. PLoS Biol 5:e94.

Strelkowa, N. and Lässig, M. 2012. Clonal interference in the evolution of influenza. Genetics 192:671–682.

Sung, W., Ackerman, M. S., Miller, S. F., Doak, T. G., and Lynch, M. 2012. Drift-barrier hypothesis and mutation-rate evolution. Proc. Natl. Acad. Sci. 109:18488–18492.

Tenaillon, O., Rodrguez-Verdugo, A., Gaut, R. L., McDonald, P., Bennett, A. F., Long, A. D., and Gaut, B. S. 2012. The molecular diversity of adaptive convergence. Science 335:457–461.

Weinreich, D. M., Delaney, N. F., DePristo, M. A., and Hartl, D. L. 2006. Darwinian evolution can follow only very few mutational paths to fitter proteins. Science 312:111–114.

Wiser, M. J., Ribeck, N., and Lenski, R. E. 2013. Long-term dynamics of adaptation in asexual populations. Science 342:1364–1367.

Wloch, D. M., Szafraniec, K., Borts, R. H., and Ko-rona, R. 2001. Direct estimate of the mutation rate and the distribution of fitness effects in the yeast Saccharomyces cerevisiae. Genetics 159:441–452.

Wright, S. 1931. Evolution in mendelian populations. Genetics 16:97–159.

Wylie, C. S. and Shakhnovich, E. I. 2011. A biophysical protein folding model accounts for most mutational fitness effects in viruses. Proceedings of the National Academy of Sciences 108:9916–9921.

Zanini, F. and Neher, R. A. 2012. Ffpopsim: An efficient forward simulation package for the evolution of large populations. Bioinformatics 28:3332–3333.

Zeyl, C. and DeVisser, J. A. G. 2001. Estimates of the rate and distribution of fitness effects of spontaneous mutation in saccharomyces cerevisiae. Genetics 157:53–61.

